# Replicative senescence as a consequence of stochastic processes

**DOI:** 10.1101/490433

**Authors:** Alejandro J. Fendrik, Lilia Romanelli, Ernesto Rotondo

**Affiliations:** Instituto de Ciencia, Universidad Nacional General Sarmiento, Buenos Aires Argentina; Consejo Nacional de Investigaciones Cientificas y Tecnicas CONICET, Buenos Aires, Argentina

## Abstract

The aging of multicellular organisms is a complex problem. It has been the subject of extensive research and have led to various theories. One of the remarkable features of aging is that is accompanied by the decrease in the rate of cell replication in tissue regeneration processes (replicative senescence). In the present work, we will show, if sporadic cell alterations occur in a homogenous set of stem cells, they lengthen the cell cycle and these are fixed with greater probability than non-altered stem cells. This effect is due to the inherent characteristics of the renewal dynamics and as time goes by it leads to a quiescence state for stem cells due to the recurrent fixation of such altered cell.

## Introduction

The effect of aging in multicellular organisms in homeostatic regime is characterized by the decline of cell renewal processes in all organs and tissues [1–5]. This diminution is closely related to the functions of stem cells (SC) responsible for cell renewal and it is more evident in the tissues of which a high rate of cell renewal, such as the skin epithelium or intestinal epithelium, is necessary. Although some variations in the behavior of SC are known with age, and studied from different the point of view, as genomic [6–8], epigenomic [9,10] and proteomic [11–13], although the scope and consequences of these changes have not been well established. Consider a tissue or part of it, whose cell renewal processes is due to small set of SC, (as in the intestinal crypts). It has been established that the population dynamics of SC in cell renewal process is neutral, from the monoclonal character of the crypts [14,15]. This was profusely studied in the framework of the dynamics of birth and death processes in a population of distinguishable but equally fit individuals, which leads to the survival of the descendants of only one of the original individual. This is a purely stochastic phenomenon and constitutes a spontaneous break of symmetry. Within the cell renewal process in the crypts a similar phenomenon occurs, because the symmetrical mitoses originating two cells involved in the differentiation process (DD division) means eliminating a SC, while symmetrical divisions originating SC (SS division) equals the birth of one of them. We show that any disturbance (from any source) sporadically affecting any of SC, as to extend their cell cycle length, leads to replicative senescence and is a consequence of the dynamics of cell renewal. Cell renewal in homeostatic regime is determined by a delicate balance between the occurrence of symmetric cell divisions of each type. So, to preserve SC population finite and avoid its extinction, the occurrence of these events must be regulated. To study this dynamic, two alternative points of view are usually adopted. In the homeostatic regime, the SC number will be considered fixed and the regulation is taken as “perfect”, that is at each symmetrical DD division it will be followed again by a symmetric SS division. This leads to the well-known Moran model [16] (widely used to study the effect of genetic drift in a small population group) and more recently, in processes of cell renewal, the fixation of neutral somatic mutations in the framework of cancer development [17]. The other point of view is to assume certain regulation for the occurrence of possible events for SC population and their daughter cells, which give rise to the complete lineage of the considered tissue. For a homogeneous population of each cellular phenotype is known as compartmental models. These models are described by a system of ODEs for the populations of each compartment [18,19]. The stationary solution of these ODEs corresponds to the average behavior of populations under homeostasis conditions. For small populations in these conditions, the result of these equations is meaningless, since the fluctuations around that average value could destabilize the population leading to extinction or exponential growth. Therefore, for these populations we must interpret ODEs as stochastic processes. If the stochastic processes lead to a stationary regime, if the ODEs and the fluctuations are neglected around the equilibrium, a Moran type model may be proposed. If an inhomogeneity is introduced in the population (as an altered SC), the process of extinction or fixation of that altered cell is necessarily stochastic and neither the ODEs nor the Moran model can be raised a priori, since the equilibrium population and its fluctuations will depend on whether or not the alteration was fixed. In what follows, we will name the alteration as a mutation, but it is not necessarily related to somatic mutation. Therefore, in the present context we use mutation in the sense of any altered cell. However there is an exception to the latter that is when the mutation was fixed but the equilibrium point or the fluctuations around it do not changed. This is the kind of mutation that will be studied in this work. This one will change the length of the cell cycle but not the balance or the regulation between the different possible events.

First we introduce a differential equation associated to the compartment of the SC and introduce the family of mutation that we will study. Then, we will calculate the fixation probabilities of these mutations by studying the associated stochastic process in two alternative ways: averaging over the realizations of the process and calculating exactly the fixed probability vector of the associated stochastic matrix. Then, since the fixation of mutations does not alter the equilibrium point and fluctuations around it, we will raise a simplified model Moran type in order to obtain a functional expression for the probability of fixation of mutations. Finally, we will show, in a simplified process to obtain functional expressions, the effect caused by the accumulation of fixations of these mutations.

## Results

### Compartment model

As mentioned in the introduction it is possible to describe population dynamics by ordinary differential equations as long as the cells of each compartment are identical. Let us consider a compartment corresponding to SCs whose populations obeys the following differential equation:

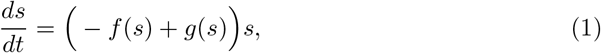

where *s* is the total number of SC present at time *t* in the compartment. The rate of disappearance and birth of SC are *f*(*s*) and *g*(*s*) respectively. If SC population remains small, this dynamics should be studied as a stochastic process and the trajectories defined by *s* = *s*(*t*), in the equilibrium regime, correspond to fluctuations around the equilibrium population given by *s*_e_ which fulfill:

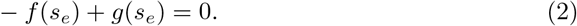

One wonders, what will happen if a different SC emerges (for example due a somatic mutation or any other alteration). Supose the desapperance (birth) rate of the mutant is *β*_1_*f*(*s*) (*β*_2_*g*(*s*)).

Clearly Eq (1) cease to have meaning because SC population in not longer homogenous. To study the resulting dynamics, we must consider it as a stochastic process of four possible events:

1. Disappearance of a “wild” stem cell (WSC).
2. Birth of a WSC.
3. Disappearance of a mutant stem cell (MSC).
4. Birth of a MSC.

The description of population dynamics through an equation such as Eq (1), will be recover if WSC or MSC are extinguish. If the latter, Eq (1) should be replaced by:

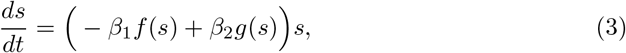

and now, the equilibrium population should be defined by:

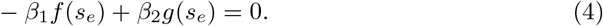

Let consider mutations where *β*_1_ = *β*_2_ = *α*. Therefore, Eq (3) is reduced to Eq (1) (it is worth to notice that there is not change in the equilibrium population) and Eq (3) is rewritten as:

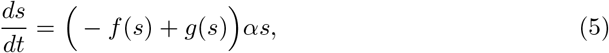

By defining *t*′ = *αt* this equation is identical to Eq (1). This means that MSCs have different length cell cycle, *τ*_*m*_, than those of WSCs, *τ*_*W*_, such that:

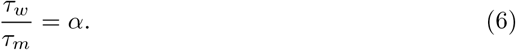

Given *α* < 1, the fixation of such mutation slows cell renewal processes, what is own aging of tissues.

In the following section we disscus the probabilities of fixing this kind of mutations.

### Fixing mutations

To analyze how mutations are fixed let consider a function *V*(*s*) as a potential:

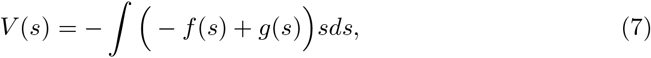

it is enough to have a deep minimum at *s* = *s*_e_ to avoid extinctions or exponential growth due to fluctuations. therefore, considering in Eq (3) we propose:

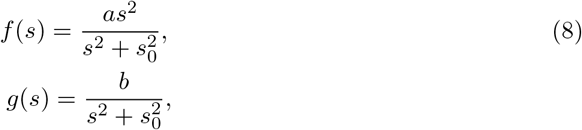

which fulfill the conditions given above. For all results shown in this paper we take *a* = 1 / day, *b* = 100 1/ day, *s*_0_ =10 and therefore *s*_e_ = 10, with these parameters the cell cycle is *τ* = 1 day for the homogenous starting system under homeostasis conditions.

We start from the state in which there are present (*s*_e_ − 1) WSCs and only one MSC, this is labeled as (*s*_e_ − 1,1). Therefore the events mentioned in the previous section are:

1. (*s*, *s*′) → (*s* − 1, *s*′),
2. (*s*, *s*′) → (*s* + 1, *s*′),
3. (*s*, *s*′) → (*s*, *s*′ − 1),
4. (*s*, *s*′) → (*s*, *s*′ + 1).

where *s* (*s*′) is the number of WSCs (MSCs). The probabilities of occurrence of these event are given in Table 1.

**Table 1.** Probabilities of the events that define the stochastic process mentioned in the text.

| Event | Probability |
| --- | --- |
| $(s, s') \rightarrow (s - 1, s')$ | $P_s^- = \frac{f(s+s')}{\mathfrak{N}} s$ |
| $(s, s') \rightarrow (s + 1, s')$ | $P_s^+ = \frac{g(s+s')}{\mathfrak{N}} s$ |
| $(s, s') \rightarrow (s, s' - 1)$ | $P_{s'}^- = \frac{f(s+s')}{\mathfrak{N}} \alpha s'$ |
| $(s, s') \rightarrow (s, s' + 1)$ | $P_{s'}^+ = \frac{g(s+s')}{\mathfrak{N}} \alpha s'$ |

Where $\mathfrak{N} = (g(s + s') + f(s + s'))(s + \alpha s')$ , $s$ ( $s'$ ) is the number of WSCs (MSCs).

We determine the probabilities of fixation by two ways:

1. Calculating the fixation frequencies over 10^6^ realizations of the stochastic process.
2. Determining the fixed vector of the stochastic matrix constructed from the probabilities shown in Table 1 (see section Methods).

The probabilities of fixation of the MSC, 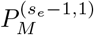, and the probabilities of fixation for one of the WSCs, 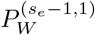, as a function of *α* are displayed in Fig 1. As expected, the curves obtained by the both method match. It can be observe when the mutation is neutral (*α* = 1) the fixation probabilities are equal for each one of SCs initially present (*s* = *s*_e_ = 10 in this case).

**Fig 1.**
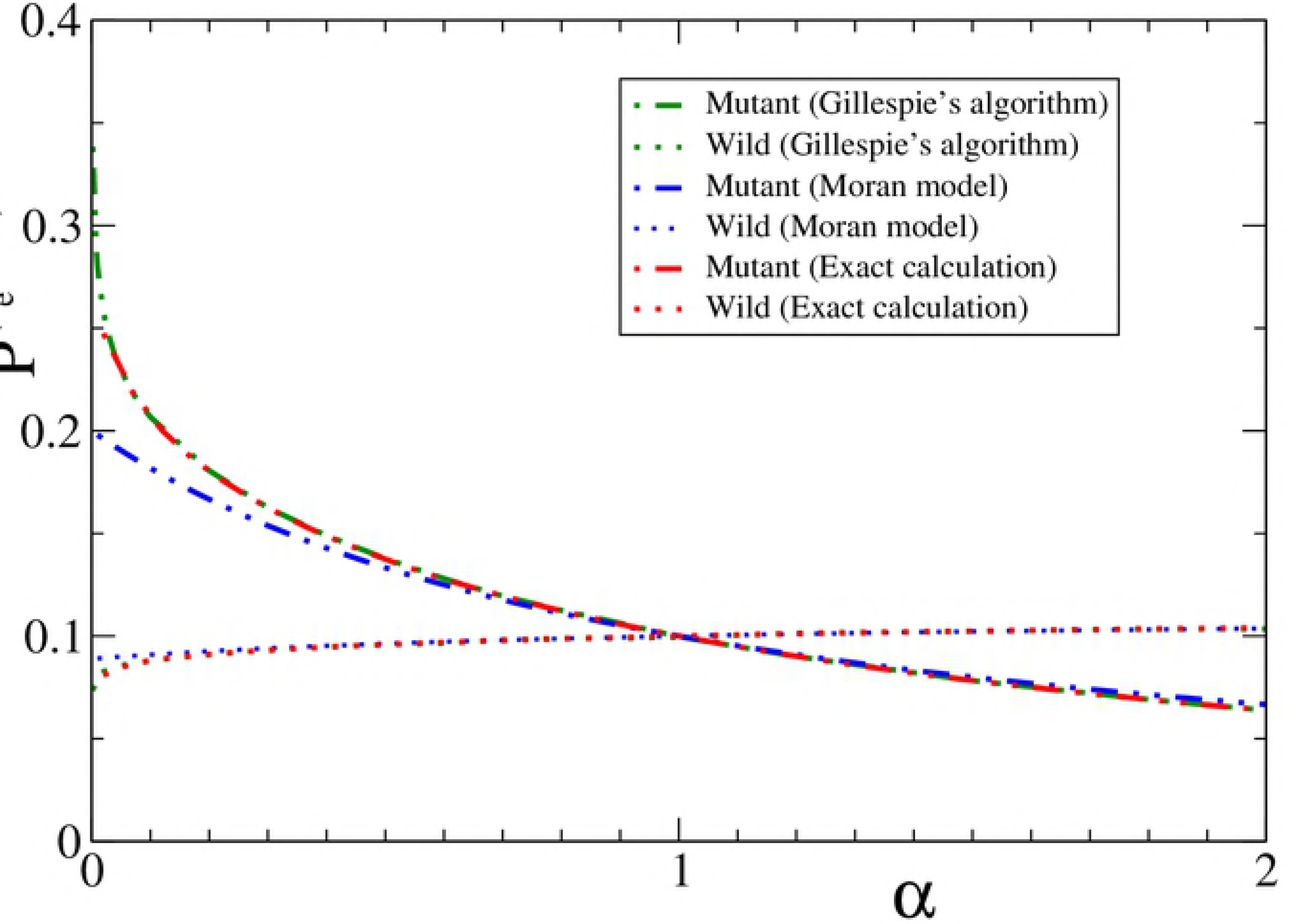
Fixation Probabilities as a function of *α*. The curves correspond to the probability of fixation of mutant and each of the WSC starting from a state ((*s*_e_ − 1), 1). The red and green curves were obtained from the stochastic process in two different ways. Those which are green correspond to the average of 10^6^ realizations of the process, while the red ones to the exact calculation of the fixed probability vector of the stochastic matrix. The blue curves correspond to considering the stochastic process as a Moran and leads to two algebraic expressions for such probabilities.

Which is striking about these results is that the probabilities of fixing of MSC increases with respect to the probability of fixing each WSC. So, in cell renewal process from symmetrical divisions, the fixing of those SC with longer cell cycle will be favored.

In next section we will propose a simplification that allow to determine the dependence of fixation probabilities on *α*.

### Simplified model

Proved stability of the population against fluctuations and since the probabilities of fixing MSCs does not modify the value of the equilibrium population (*s*_e_) nor the fluctuation around it, we can simplify the problem by keeping the total number of SC fixed (*s* + *s*′ = *s*_e_).

Considering now the stochastic process as joint event of disapearances and births, the total number of SC remains constant. Now the possible states are characterized by (*s*_e_ − *s*′) for WSC and *s*′ for MSC present. Therefore, the stochastic process includes only the following events:

1. (*s*_e_ − *s*′, *s*′) → (*s*_e_ − *s*′ − 1, *s*′ + 1),
2. (*s*_e_ − *s*′, *s*′) → (*s*_e_ − *s*′ + 1, *s*′ − 1),
3. (*s*_e_ − *s*′, *s*′) → (*s*_e_ − *s*′, *s*′).

The associated probabilities are shown in Table 2.

**Table 2.** Probabilities of the events that define the stochastic process mentioned in the text.

| Event | Probability |
| --- | --- |
| $(s_e - s', s') \rightarrow (s_e - s' - 1, s' + 1)$ | $\frac{(s_e - s')}{(s_e - s' + \alpha s')} \frac{\alpha s'}{(s_e - s' - 1 + \alpha s')}$ |
| $(s_e - s', s') \rightarrow (s_e - s' + 1, s' - 1)$ | $\frac{\alpha s'}{(s_e - s' + \alpha s')} \frac{(s_e - s')}{(s_e - s' + \alpha(s' - 1))}$ |
| $(s_e - s', s') \rightarrow (s_e - s', s')$ | $\frac{(s_e - s')}{(s_e - s' + \alpha s')} \frac{(s_e - s' - 1)}{(s_e - s' - 1 + \alpha s')} + \frac{\alpha s'}{(s_e - s' + \alpha s')} \frac{(s' - 1)\alpha}{(s_e - s' + \alpha(s' - 1))}$ |

The resulting stochastic matrix has two absorbent states: (*s*_e_, 0) and (0, *s*_e_) (all WSCs or MSC). Starting from the initial state (*s*_e_ − *s*′, *s*′), the probability of ending at (0, *s*_e_) for *α* > 0 results (see see section Methods):

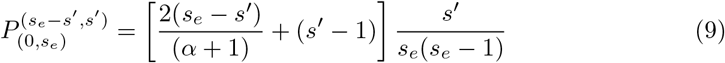

and the probability of ending at (0, *s*_e_) is its complementary:

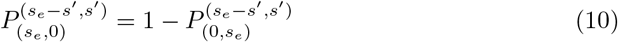

The latter correspond to a not fixed mutation but any of (*s*_e_ − 1) WSCs, while the probability of one of them is fixed is:

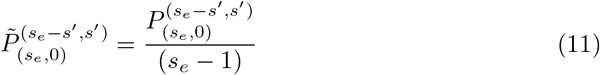

We are interested in studiying what happen when within a population of identical cells a mutation of one of them occurs. Starting from (*s*_e_ − 1,1):

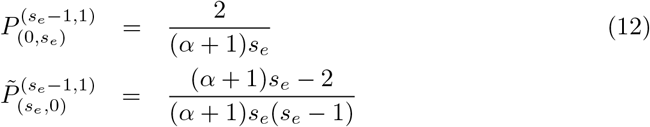

These probabilities should be compared with those that we have obtained in the precedent section 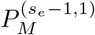 and 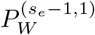.

Fixation probability of MSC leads to 1/*s*_e_ if the mutation is neutral and it grows (decreases) for *α* less (greater) than one. Figure 1 shows the curves obtained from Eqs (12) which agrees to those calculated in the previous section for a significant range of *α* values.

Let us remark regarding the simplified model, it becomes unrealistic if the mutation changes the balance between births and disappearances (changes in *β*_1_ and *β*_2_) since the fixation of mutant not only changes the equilibrium population but could also lead to system instability.

### Fixation of successive mutations and their effect

Slow SC fixation are the effect of reducing the rate in cell renewal processes. This decrease should not necessary occurs as the result of discrete events affecting cell cycle length, such as mutation. Indeed, the cell cycle due to WSC follows a distribution around some mean value and the effect shown in previous sections tends to fix more the SCs whose cell cycle length are less than the average.

Now let us consider the occurrence of successive mutations and their effect on cell cycle length. Currently there is not available experimental data of these possible mutations since they are not trivially detectable except for the observed effect in the slowness of celular renewal dynamics (as the aging process). Given the occurrence of *m* mutation per year on average, the process of successive mutations generate the tree diagram shown in Fig 2. We assume that the effect of each mutation is the same for SC population not previously fixed, that is to say the effect is always vary the cell cycle length in the same percentage *α* = *τ*_*w*_/*τ*_*m*_. Therefore, 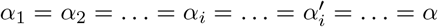.

**Fig 2.**
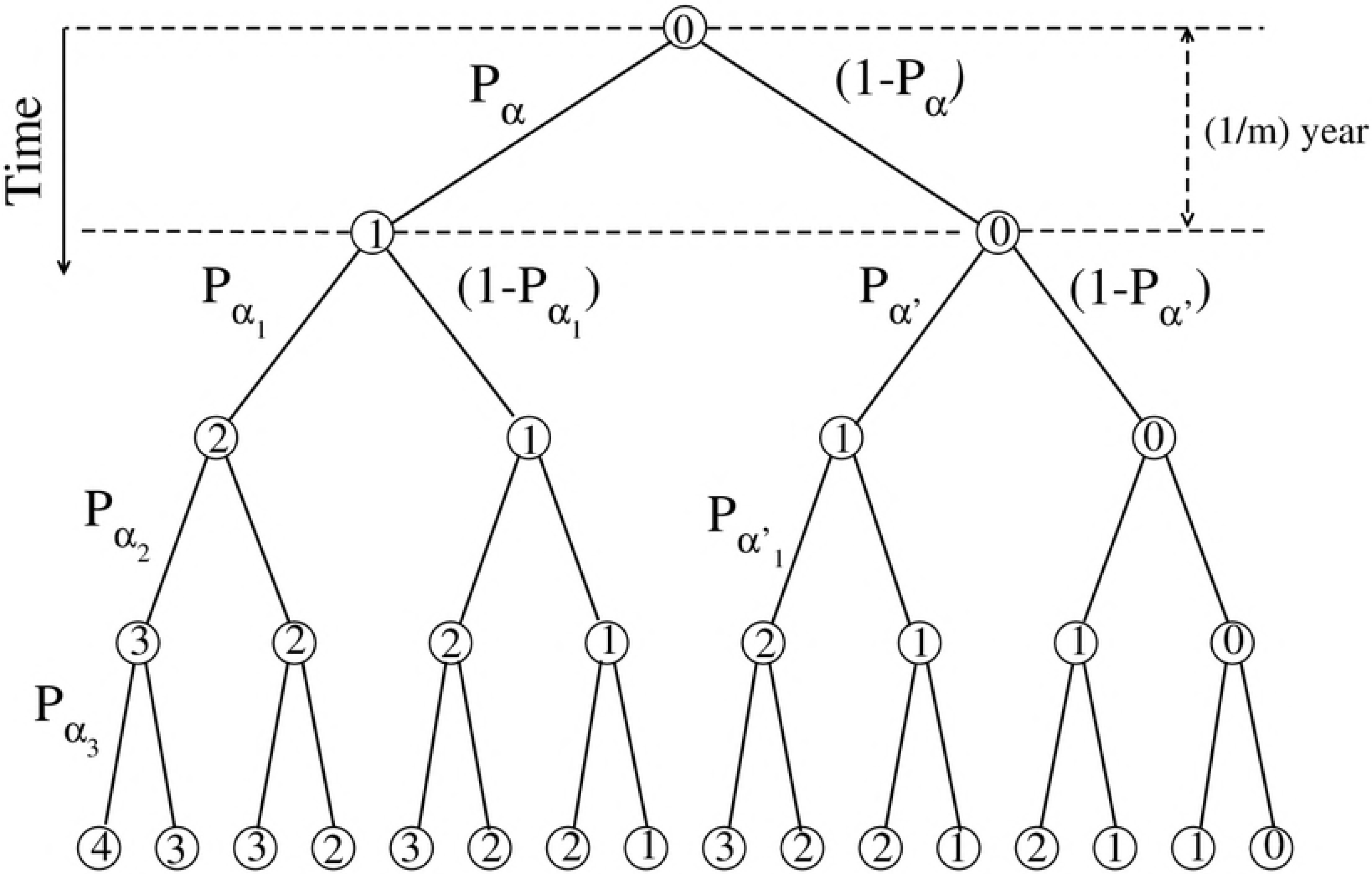
Tree Diagram for successive mutations. The diagram illustrates the possible final states of the system after four mutations, assuming that they occur on average, m annual mutations. The numbers indicate the number of mutations that were fixed.

So, after *k* fixed mutations the cell cycle length will be *α*^*k*^ = 1/*τ* (we assume as a unit the cell cycle length of the original population).

The probability that of *N* mutations *k* are fixed is:

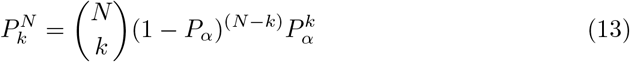

where *P*_*α*_ is the probability of fixation of one mutation. Thus, the cell cycle length after *N* mutation is:

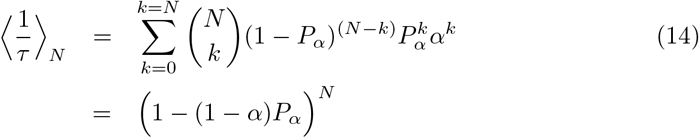

and taking into account the first of Eqs (12):

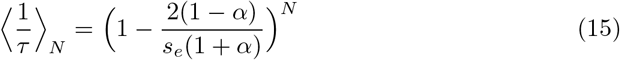

It is possible generalize these results for more than one *α*. If there are two different mutations characterizes by *α* and *α*′, and assuming that the mutation associated with *α* occurs with probability *p*_+_ and *p*_−_ for *α*′, (*p*_+_ + *p*_−_ = 1), we obtain:

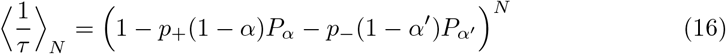

We will now assume that *α* increases the cell cycle length to the same extend that *α*′ decreases it, this means *α* + *α*′ = 2. Therefore we have:

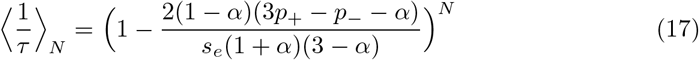

Figure 3 shows the results of this last expression as function of α for several probabilities of occurrence *p*_+_ and *p*_−_ after 200 mutations assuming *m* = 2. When sporadic mutations only lengthen the cell cycle (*p*_+_ = 1, *p*_−_ = 0), the average duration of that cycle increases as α decreases (recall α is the exchange value in the cycle for a sporadic mutation) for *α* ≤ 0.5 the average state of the CS is of total quiescence, while if *α* = 0.8 the average cell cycle after 100 years is ten times larger than a younger system. By including the probability of a mutation that decreases the cell cycle (*p*_−_ ≠ 0), the effect of decreasing α is to increase the cell cycle even if the increases are more likely than decreases, if α is small enough

**Fig 3.**
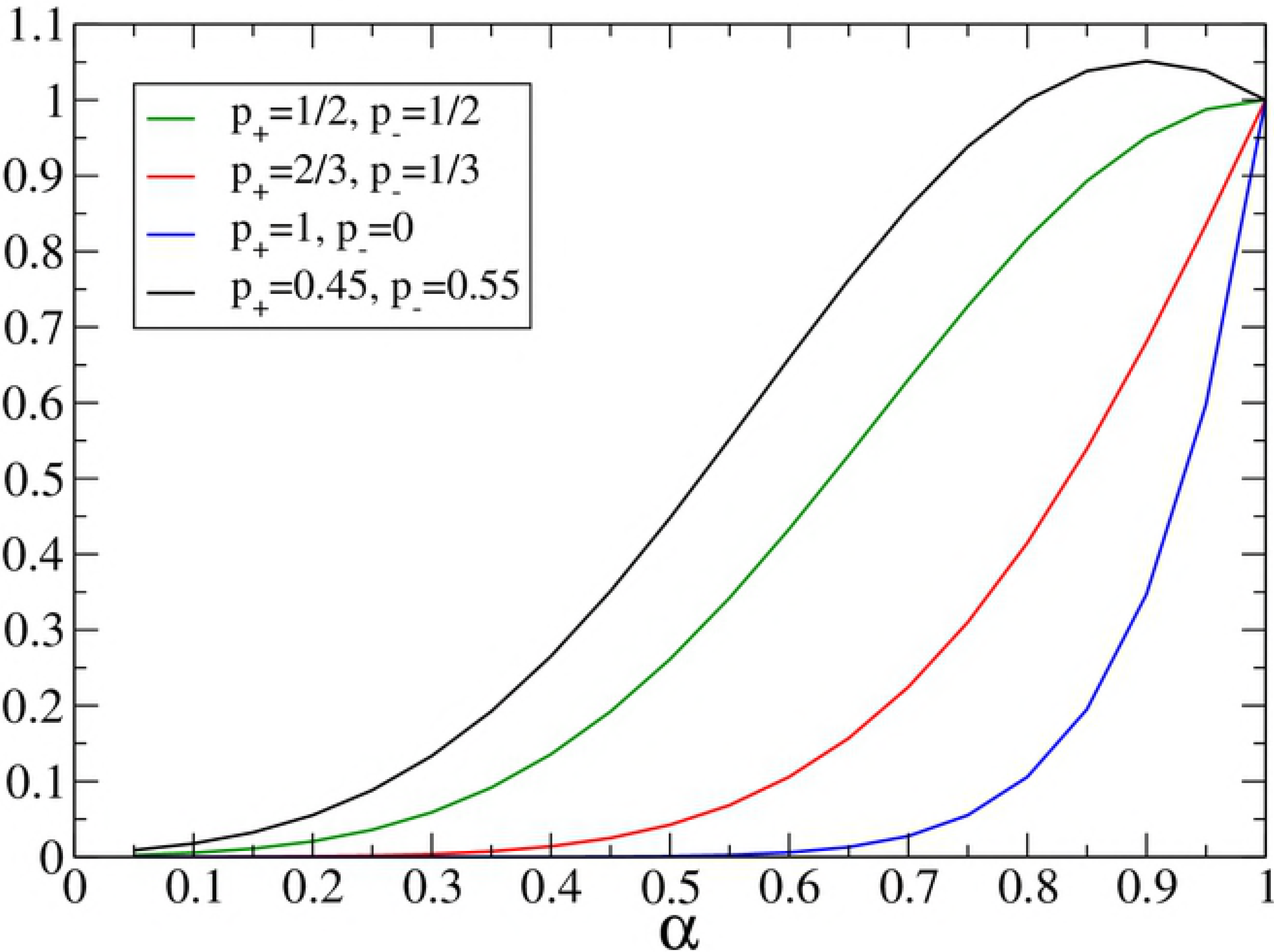
Length of the cell cycle as a function of *α* after 200 mutations. The graph illustrates the effects in the cell cycle *τ* after 200 mutations assuming that there may be two kind of mutations. One of them increases the duration of *τ* (*α* < 1) and decreases it with probability *p*_+_, while the other decreases it (*α*′ > 1) with probability *p*_−_ and *α* + *α*′ = 2 (see Eq 17). The blue one corresponds to *p*_+_ = 1, *p*_−_ = 0, for the mutation which increases the cell cycle. The curvesred, green and black to *p*_+_ = 2/3, *p*_−_ = 1/3, *p*_+_ = 1/2, *p*_−_ = 1/2 and *p*_+_ = 0.45, *p*_−_ = 0.55.

## Methods

The results shown above were found by numerical calculation and analytical derivations.

### Numerical calculations

These were carried out in two ways.

1. To simulate the stochastic process by using Gillespie′s Algorithm [20].
2. Detrmination of the fixed probability vector associated to stochastic matrix. The stochastic matrix is:

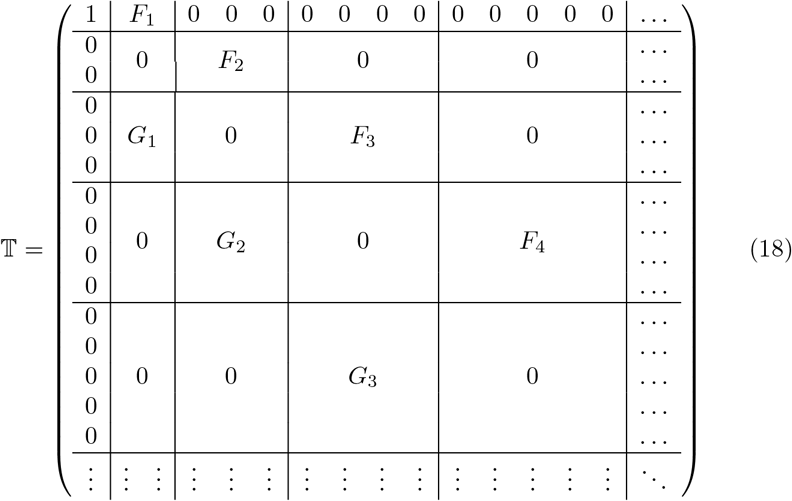

where *F*_*M*_ and *G*_*M*_ are *M* × (*M* + 1) and (*M* + 2) × (*M* + 1) matrices respectively. Their elements are:

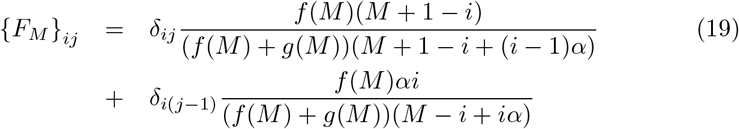

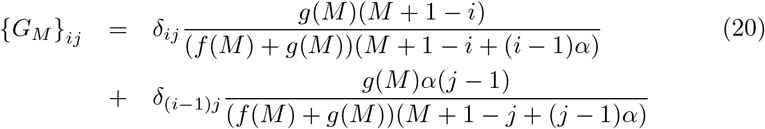

The matrix 𝕋 acting on the probability vector **p**:

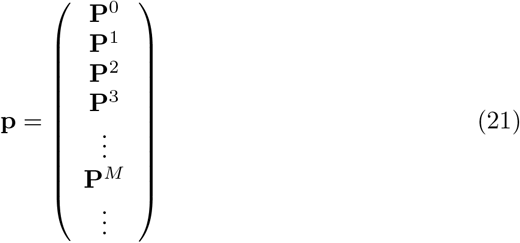

where:

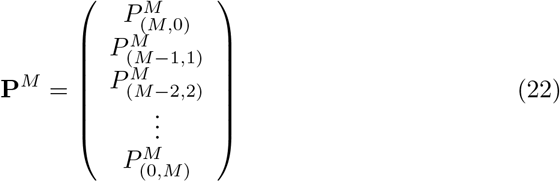

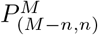 is the probability that the system has (*M* − *n*) WSCs and *n* MSCs. The stochastic process changes the total number of SCs. In fact, each event of the process creates or destroys a cell. On the other hand, the matrix 𝕋 connects even states (*M* even) with odd (odd *M*) or odd with even ones. Taking advantage of this fact we can reduce 𝕋. Rearranging **p** as:

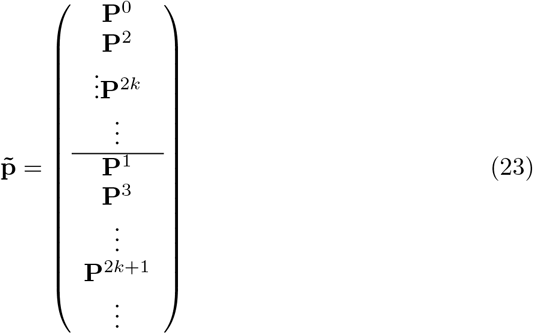

We obtain:

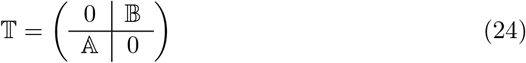

where:

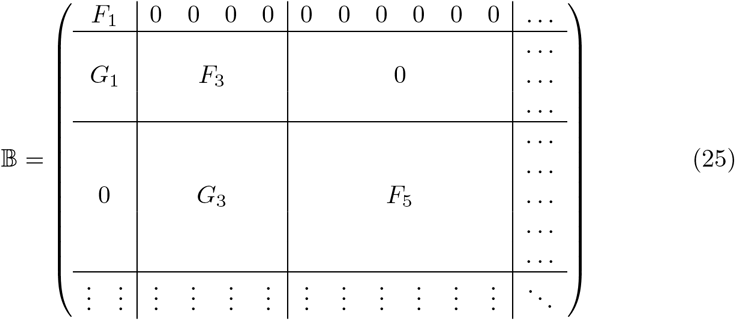

and:

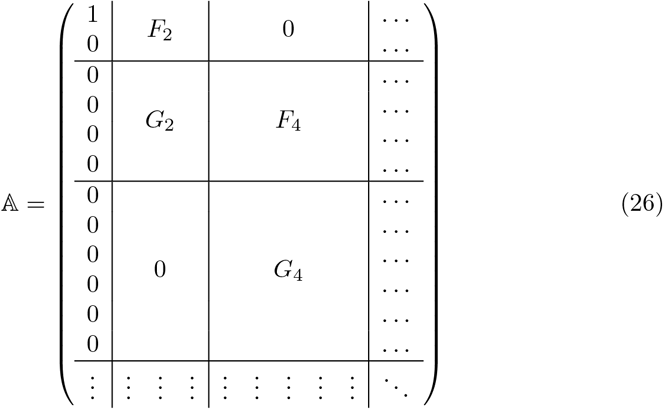

Therefore 𝕋^2^ has one representation on the even space (𝔹𝔸) and other on the odd space (𝔸𝔹).

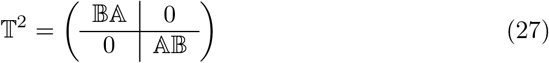

To perform calculations with these expressions, it must be taken into account that the matrix T (like A and B) has infinite dimensions. So, they must be truncated. This truncation is not always possible because we must guaranteed that after it the matrices remain stochastic, then it is necessary that:

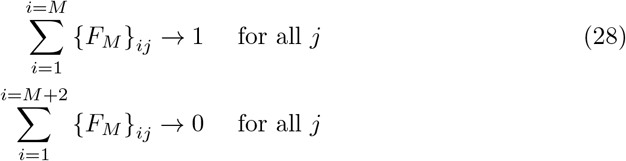

for some *M*. This will depend on the behavior of *f*(*M*) and *g*(*M*).

For the result shown in Fig 1, we obtain the steady state probability vector **p̃**_e_ starting from the probability vector with 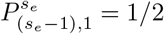, 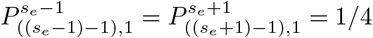 where *s*_e_ satisfies Eq (2).

The non-zero components of **p̃**_e_ are 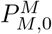 and 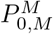 corresponding to fixation probabilities of WSCs or to fixation of *MSC* respectively. Therefore:

- The probabilities of find the system with *N* MSC in homeostatic conditions are:

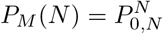
- The probabilities of find the system with *N* WSC are:

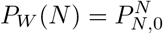
- The probabilities of find the system with *N* WSCs descendent of one of the SCs, initially present are:

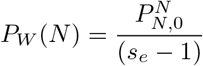
- The probabilities to find the system with *N* SCs in homeostatic conditions are:

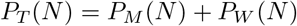

Such distribution are displayed en Fig 4.

**Fig 4.**
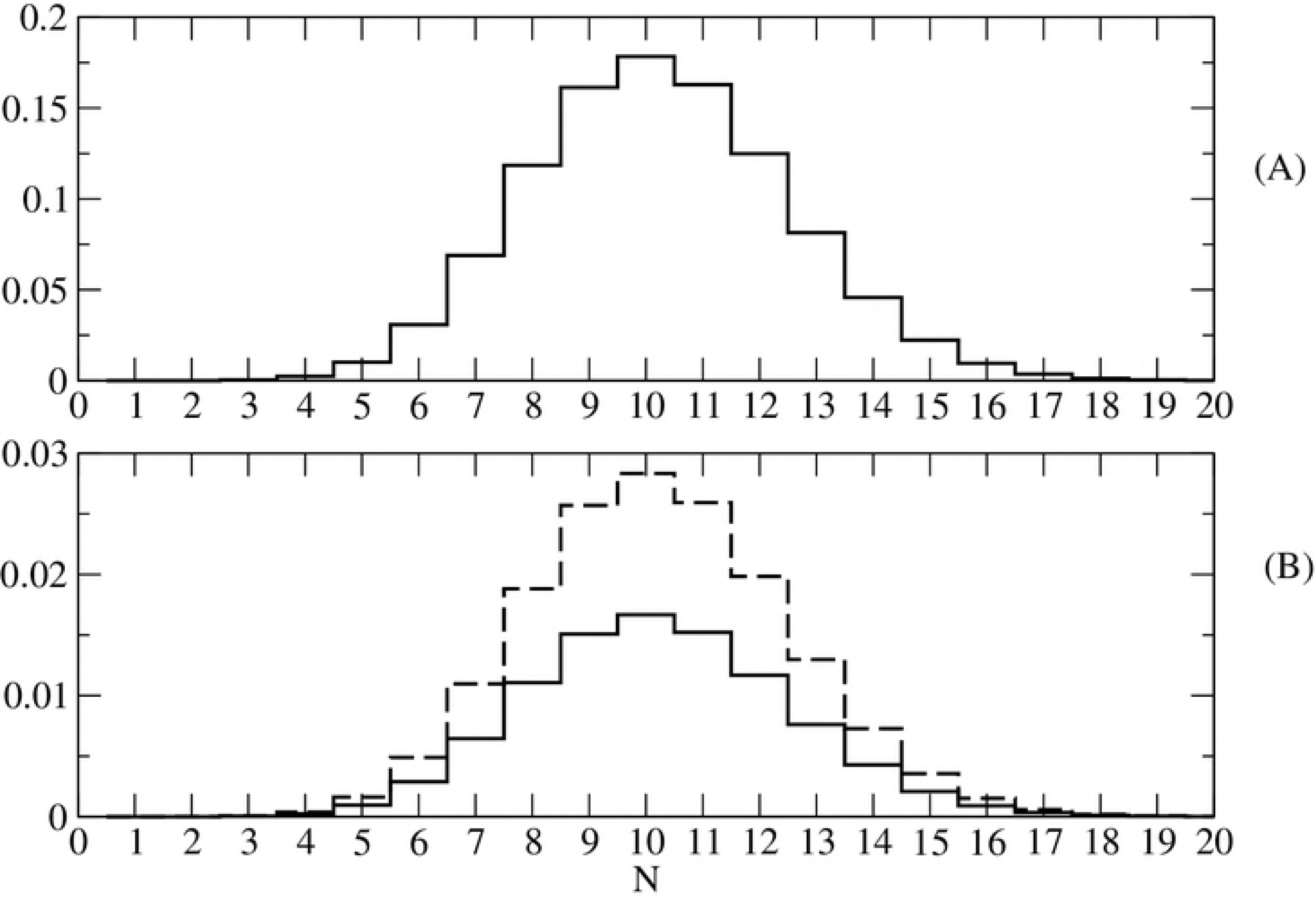
P(N) vs. N. A) Probabilities to find the system with *N* SCs in homeostatic conditions. B) The probabilities to find the system with *N* MSCs (dashed line) and the probabilities to find the system with N WSCs descendent of one of the SCs, initially present (full line) for *α* = 0.3.

On the other hand:

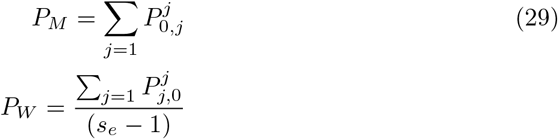

These probabilities as a function of a are shown in Fig 1 of the paper.

### Analytical derivations

Regarding simplified model, we will show how obtain the probabilities of fixation of MSC starting from a given state (*N* − *n*,*n*) (that is (*N* − *n*) WSC, *n* MSC). The stochastic matrix 𝕋 acts on column vectors like:

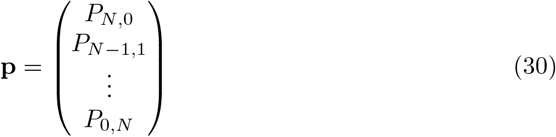

where *P*_*N−n.n*_ is the probability that the system is in the state (*N* − *n*, *n*).

The matrix 𝕋 is tridiagonal and its elements are:

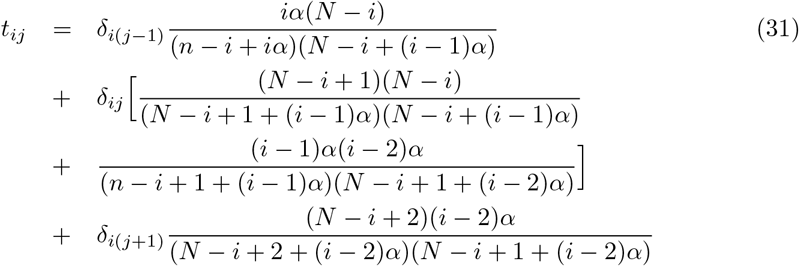

for 2 ≤ *i*, *j* ≤ *N* and

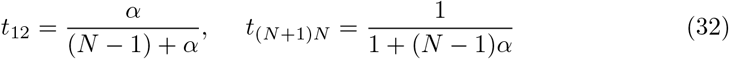

𝕋 can be write as:

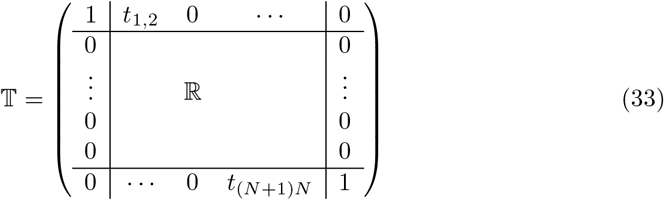

where {ℝ}_*ij*_ = *t*_(*i*−1)(*j*−1)_.

Defining the row vectors **v** and **u** as

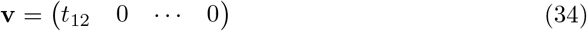

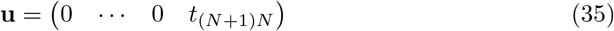

results

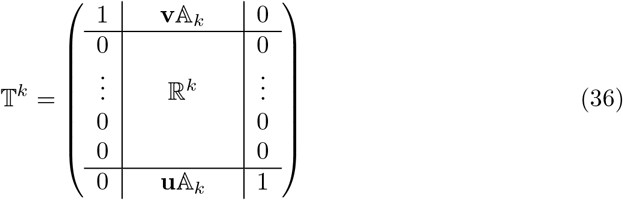

where

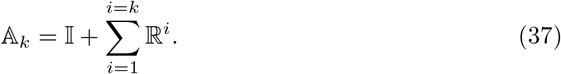

And due to

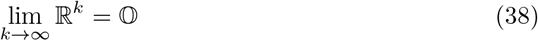

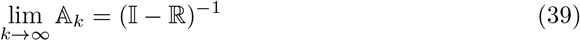

We obtain:

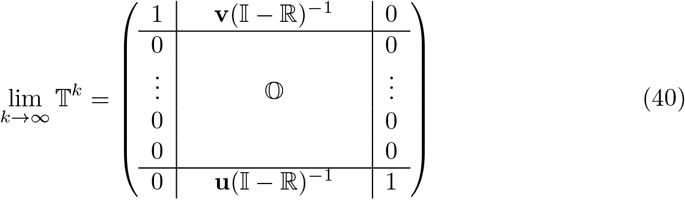

The last row of the latter (0, **u**(𝕀 − ℝ)^−1^,1) contains the probabilities of fixing MSC. In fact,due to the vector **u** it is not necessary to calculate the whole (𝕀 − ℝ)^−1^. We only need to know the last row.

In this way we obtain:

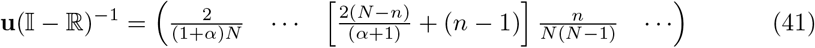

## Discussion and Conclusion

Whereas cell renewal follows a neutral dynamic from a small set of SC, we have shown that fixation SC with longer cell cycle is favored and therefore, as time goes by, such renewal is becoming slower. This does not mean that cellular senescence is due to this effect (it is due purely to the characteristics of renewal dynamics). However, and admitting that the causes of replicative senescence could be eliminated, the intrinsic dynamics itself leads to the inevitable slowing down of the rate of cellular renewal, to the point of being meaningless from a certain chronological age. We have also shown that the most probable effect is the fixation of the slow SC, from those SC affecting only the cell cycle length. As cellular cycles of an identical set of SCs obeys a distribution, (not a fix values) it is expected that the effect acts continuously favoring the fixation of the SC whose cell cycle falls in the region of lower cellular cycles of such distribution. On the other hand, we have assumed cell renewal occurs from symmetrical divisions of SCs. The asymmetric divisions that actually exist [21,22] do not vary the balance expressed by Eq(1) corresponding to SC compartment. (although they do it in the following compartments corresponding to cells already committed in the process of differentiation). From this point of view the inclusion of asymmetric divisions leads to the fact that not all mitosis participate in the dynamics described above and therefore their presence drive to an effective lengthening of the cell cycle. This means that possible mutations that modify the balance between symmetric and asymmetric divisions in favour of the latter would be more likely to be fixed.

## Acknowledgments

We are in debt to A. Braunstein and M.V.Reale for helpful discussions. AJF and LR thanks the hospitality of the Italian Institute of Genomic Medicine at Torino. The authors acknowledge funding from EU, Horizon 2020, Marie Sklodowska - Curie grant agreementNo 734439 (INFERNET)

